# Sepal shape variability is robust to cell size heterogeneity in Arabidopsis

**DOI:** 10.1101/2024.04.04.588049

**Authors:** Duy-Chi Trinh, Claire Lionnet, Christophe Trehin, Olivier Hamant

## Abstract

How organisms produce organs with robust shapes and sizes is still an open question. In recent years, the Arabidopsis sepal has been used as a model system to study this question because of its highly reproducible shape and size. One interesting aspect of the sepal is that its epidermis contains cells with very different sizes. Previous reports had qualitatively shown that sepals with more or less giant cells exhibit comparable final size and shape. Here we investigate this question using quantitative approaches. We find that a mixed population of cell size modestly contribute to the normal width of the sepal, but is not essential for its shape robustness. Furthermore, in a mutant with increased cell and organ growth variability, the change in final sepal shape caused by giant cells is exaggerated, but the shape robustness is not affected. This formally demonstrates that sepal shape variability is robust to cell size heterogeneity.

**Main conclusion:** A mixed population of cells with varied sizes plays a limited role in ensuring the symmetrical shape of the sepal, and is not essential for sepal shape robustness in Arabidopsis.

## Introduction

How organisms produce organs with robust shapes and sizes is one of the central mysteries of development in all kingdoms [1]. Cell size can have an ambiguous contribution to organ shape. There are examples where increasing cell size also increases organ shape, and others where increase in cell size is compensated (e.g. by decreasing cell number) [2,3]. In either case, how this affects the standard deviation in organ shape (a proxy for organ shape robustness) is ill-documented. This is what we study here.

The Arabidopsis sepal, the protective organ of a flower, offers an excellent model to answer that question because of its highly reproducible shape and size and easy access, among other reasons [4]. The sepal initiates from a flower meristem, and quickly grows following a well-documented pattern of cell growth and division [5–7].

Using this model, several possible mechanisms for shape robustness have been put forward. These mechanisms may be at the whole organ scale, such as timing of initiation and growth arrest [5,7]. They may also involve activities at the cellular scale, such as the constant reorientation of cell growth direction to achieve a robust average one, or the mechanical shielding of cells neighboring fast-growing ones [8,9]. Recently, we investigated the possible impacts of increased transcriptional noise to shape robustness using a mutant of VERNALIZATION INDEPENDENCE 3 (VIP3), a subunit of the Polymerase-associated factor 1 complex (Paf1C). VIP3 interacts with other subunits of Paf1C to control transcription of multiple genes [10–12]. In the *vip3-1* mutant, transcriptional noise and growth rates between neighboring cells are more variable compared to the WT [6]. This increased local growth heterogeneity interferes with the formation of the large-scale growth pattern typically seen in the WT sepals, manifested as a delayed growth arrest at the sepal tip of the *vip3* mutant [5,6].

The epidermal layer of a sepal consists of cells of vastly different size, which are usually divided into two different cell types: smaller cells and giant cells. Smaller cells are the product of frequent cell division, while giant cells result from early termination of cell division and subsequent endoduplication [4,13,14]. Giant cells are very long cells that can extend from the base to the tip of the sepal, and they are usually quite straight [15,16]. Despite the variability in cell sizes and growth rates, all cells experience a similar sigmoid curve where growth is initially slow, then accelerate, and then slow down again, and they all reach the same maximum of growth rate, albeit at different times [17]. Past studies have shown that the sepals in the wild type and mutants do not differ majorly, whether the mutant epidermis lacks giant cells or instead, is made up of giant cells only [13,18]. However, the aspect of shape robustness was not characterized quantitatively, hence whether more subtle effects are induced when the ratio of giant vs. small cells is affected remains to be investigated. In the present study, we examined the effects of a dominantly giant cell population to sepal shape robustness in the wild type background,, also testing the response in the *vip3-1* mutant background where mechanisms for shape robustness are compromised.

## MATERIALS AND METHODS

### Plant materials and growth conditions

All experiments were performed on Col-0 ecotype. The *vip3-1* (Salk_139885) mutant is described in [19], and the *pATML1::LGO* line in [13]. The double mutant *vip3-1 pATML1::LGO* was generated by crossing, with *pATML1::LGO* being the male parent plant.

Plants were grown on soil 20°C in short-day conditions (8h light/16h dark) for 3 weeks then transferred to long-day conditions (16h light/8h dark cycle).

### Sepal parameter measurements

To compare sepals of different phenotypes, mature flowers of stage 14 as described in [20] were used. The abaxial sepals were removed from the flowers and placed as flat as possible on double-sided tape on a microscope slide, over a black background. The images were taken with a Leica binocular equipped with a camera. To extract sepal contours and morphological parameters such as length, width and aspect ratio, the program called SepalContour was used as originally described in [8].

### Quantification of sepal shape variability

To quantify shape variability from the sepal contours extracted by the SepalContour tool, another program called Contour Analysis also originally described in [8] was used. For a given genotype, the contours of all abaxial sepals were normalized to their area, and an average contour is calculated. The squared deviation of each contour from the average contour is then calculated (S_2_ value). These S_2_ values were used to report shape variability of sepals (higher values mean more variable shape).

### The maximal width position of the sepal

The maximal width position (MWP) of the sepal is defined as the position along the sepal where its width is largest. To identify this position, an ImageJ macro was written to scan along the sepal contour (a product of the SepalContour tool) and identify the sepal length as well as the maximal width position. This position is relative to the length of the sepal and is expressed in percentage.

### SEM images of sepals

Scanning electron microscope (SEM) images of sepals were taken with a HIROX SH3000 tabletop microscope. The chamber was cooled down to -20°C for at least 30min in advance. Sepals or floral buds were fixed on the conductive carbon double-sided tape (Nisshin EM Co., Ltd, TOKYO) on a holder, and then introduced into the chamber. After sealing the chamber, it was vacuumed and a current of 5kV was applied. The imaging was done at the magnification of 70X or 100X, using default settings.

## RESULTS AND DISCUSSION

To check whether a mix population of small and giant cells seen in the wild-type sepal is essential for its shape, we used the transgenic line *pATML1::LGO* where epidermal cells experience endoreduplication to become giant cells. *LGO (LOSS OF GIANT CELLS FROM ORGANS)* encodes a cyclin-dependent kinase inhibitor, while the *ATML1 (MERISTEM LAYER 1)* promoter drives the expression specifically in the epidermis. While a wild-type sepal exhibits cells of various sizes, those expressing *pATML1::LGO* produce long giant cells which make up most of the epidermal cell population (Figure 1A) [21]. To analyze the effects of giant cell proportion on organ shape and shape variability, we used the SepalContour tool described in [8] to extract the contour as well as to measure the aspect ratio (width/length) of each sepal (Figure 1B-C; lengths and widths in mm are provided in Supplemental Table 1). From all individual contours, an average contour and a score of shape variability (*S*_*2*_) are calculated [6,8]. The average contour means the average shape of the sepals in a given genotype. The shape variability tells us if individual sepals have similar or different shapes.

**Figure 1.**
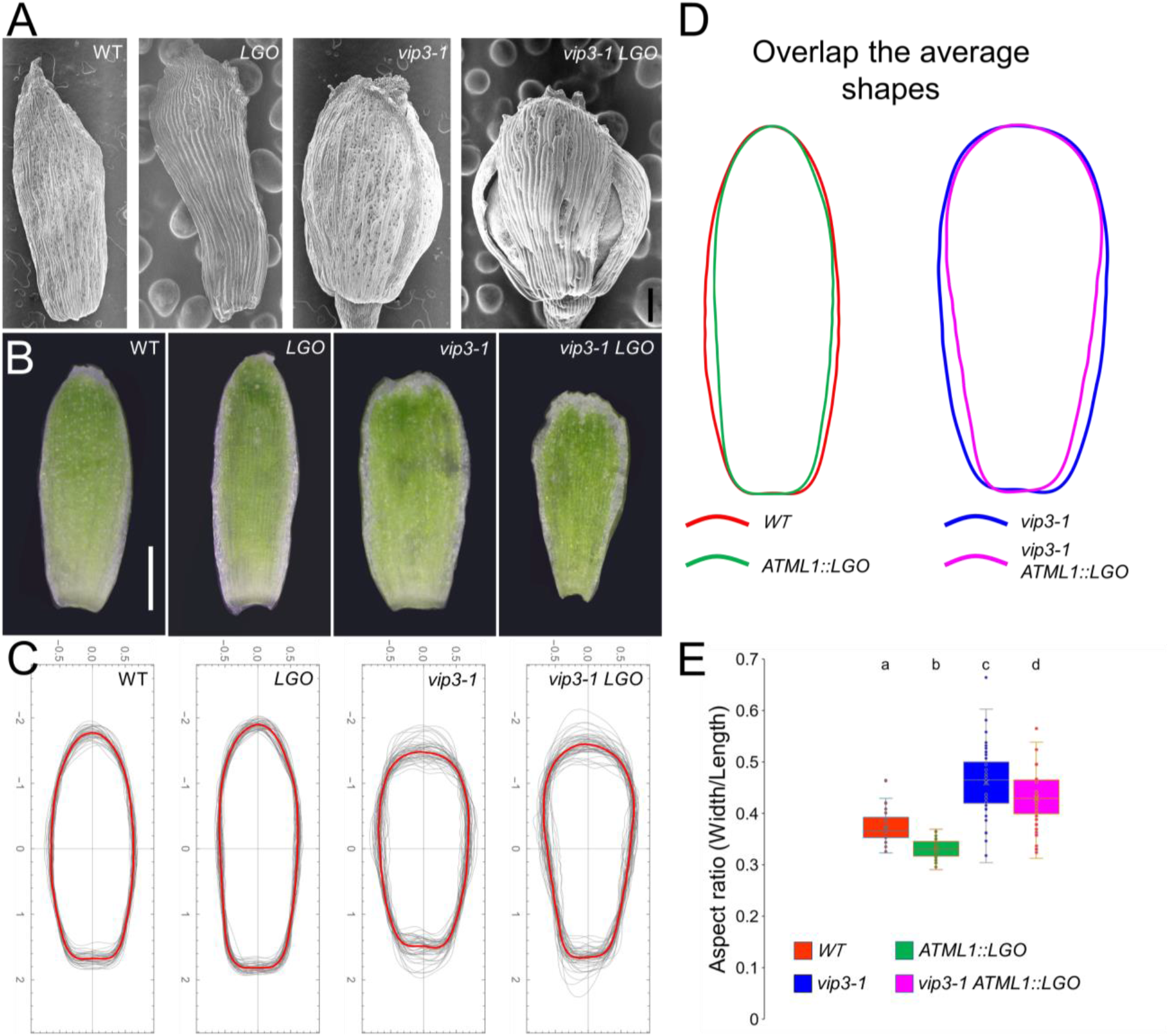
The effects of giant cells on sepal shape (A) Scanning electron microscopy pictures of wild-type, *pATML1::LGO (LGO), vip3-1* and *vip3-1 pATML1::LGO (vip3-1 LGO)* sepals. Giant cells make up most of the outer epidermal cell population in *LGO* and *vip3-1 LGO* sepals. n ≥ 5 samples for each genotype. Scale bar = 200μm. (B) Representative images of mature wild-type, *pATML1::LGO (LGO), vip3-1* and *vip3-1 pATML1::LGO (vip3-1 LGO)* sepals. Note the similarity between sepals in (B) and the corresponding average sepal contours in (C). Scale bar = 0.5mm. (C) Plots showing the contours of sepals of the four genotypes. The contours are normalized to the area. The red outlines are the average shapes. n = 40, 48, 50, 53 sepals for wild type, *pATML1::LGO (LGO), vip3-1* and *vip3-1 LGO*, respectively. (D) Overlapping average shapes of wild type and *pATML1::LGO* (upper half), and of *vip3-1* and *vip3-1 LGO* (lower half). (E) Aspect ratios (width/length) of sepals of the four genotypes. A higher aspect ratio means a wider shape, and *vice versa*. n = 40, 48, 50, 53 sepals for wild type, *pATML1::LGO (LGO), vip3-1* and *vip3-1 LGO*, respectively. Welch’s t-test. Different letters indicate statistically significant differences (*p* ≤ 0.05 for *vip3-1* and *vip3-1 pATML1::LGO* comparison, *p* ≤ 0.0001 for other pairwise comparisons).

To better see the change in the sepal shape, we overlapped the average contour of wild-type and *pATML1::LGO* sepals, and found that despite their vastly different cell populations, they are in fact very similar in shape (Figure 1D, left). This is consistent with previous qualitative observations [13,18].

We also found that *pATML1::LGO* sepals were slightly narrower, when compared to the wild type, as evidenced by a lower aspect ratio (0.37 for wild type and 0.33 for *pATML1::LGO*, Figure 1E). The narrowing of the *pATML1::LGO* sepals was more pronounced near the middle of the sepal. To characterize this change, we identified the maximal width position (MWP) of the sepal, which is the position along the sepal where its width is largest. A score greater than 0.5 means that the sepal is widest at a position closer to the sepal base, while a MWP smaller than 0.5 means that the sepal is widest closer to the tip. This MWP index can distinguish between two shapes of the same aspect ratio and is a potentially useful morphological parameter (Figure 2A). An ImageJ macro was written to scan along the sepal contour (a product of the SepalContour tool) and identify the sepal length as well as the MWP. Using this index, we found that the wild type produces symmetric sepals with the MWP around the middle. Consistent with what we noticed, the MWP index of the *pATML1::*LGO line is lower than that of the wild type (MWP_WT_ = 0.50, MWP_*pATML1::LGO*_ = 0.41, Figure 1G), meaning that the MWP of *pATML1::LGO* sepals is in the middle, closer to the tip. Because these modifications remain minor, this analysis rather confirms that sepal shape is robust to cell size perturbation.

**Figure 2.**
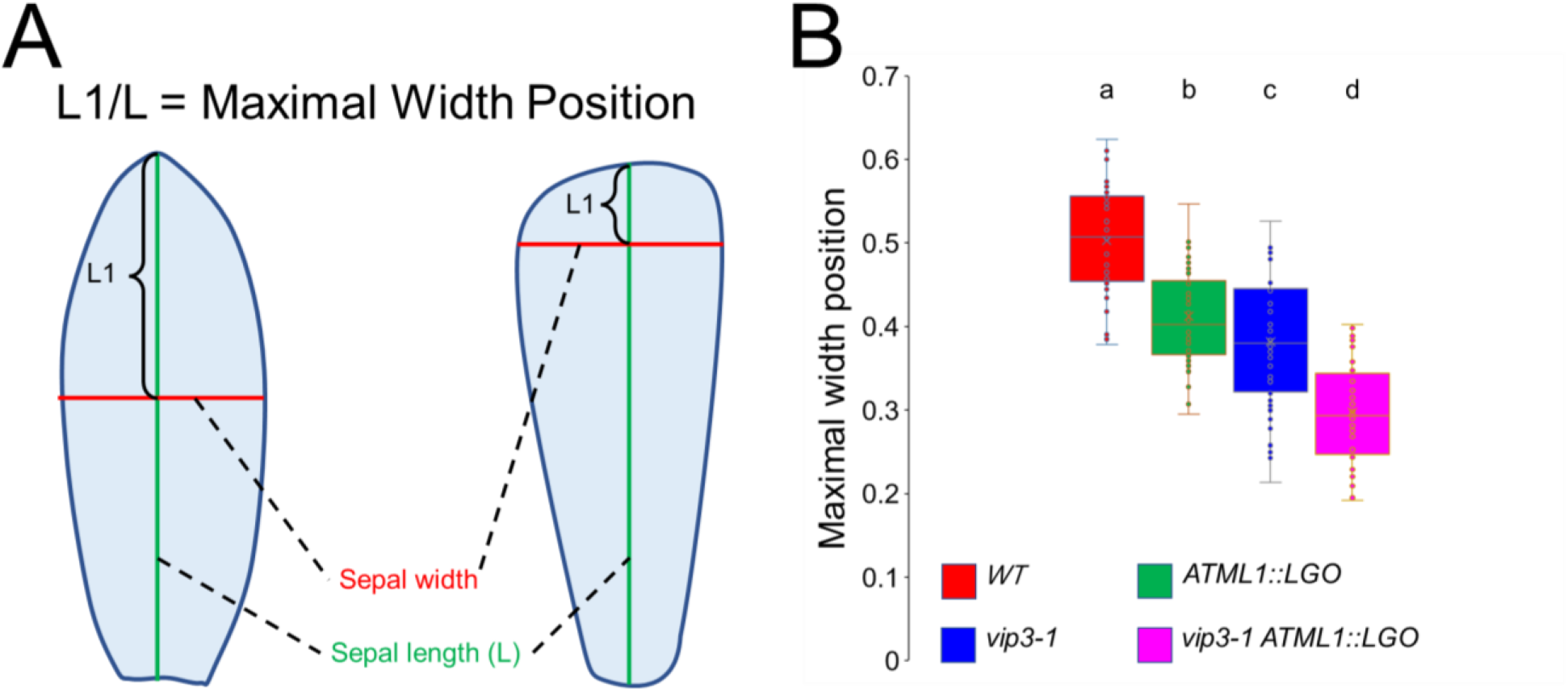
The Maximal width position index to quantify the effects of giant cells on sepal shape (A) Two shapes of the same aspect ratio can be vastly different. To distinguish them, we identify the Maximal width position (MWP) where the width is widest along the sepal length. The ratio L1/L is the MWP index, with L being the sepal length, and L1 being the distance from the tip to the MWP. (B) Maximal width position (MWP) index of sepals of the four genotypes. A lower MWP index means the sepal is widest near the tip. n = 40, 48, 50, 53 sepals for wild type, *pATML1::LGO (LGO), vip3-1* and *vip3-1 LGO*, respectively. 2-sided Welch’s t-test. Different letters indicate statistically significant differences (*p* ≤ 0.05 for *pATML1::LGO* and *vip3-1* comparison, *p* ≤ 0.0001 for other pairwise comparisons).

Yet similar shape averages do not necessarily mean similar standard deviation. To check whether more giant cells in sepals make them more or less variable in shape, we calculated shape variability (S_2_ score expressing the squared deviation of sepal contours from the average contour) using the SepalContour tool. We found that wild type and *pATML1::LGO* sepals essentially have the same level of variability (*S*_*2*_ score as median ± SE = 1.21 ± 0.29 10^−3^ for wild type, = 1.52 ± 0.15 10^−3^ for *pATML1::LGO*) (Figure 3). The data shows that sepals with only one type of cells (giant cells) can still exhibit wild-type-level shape variability.

**Figure 3.**
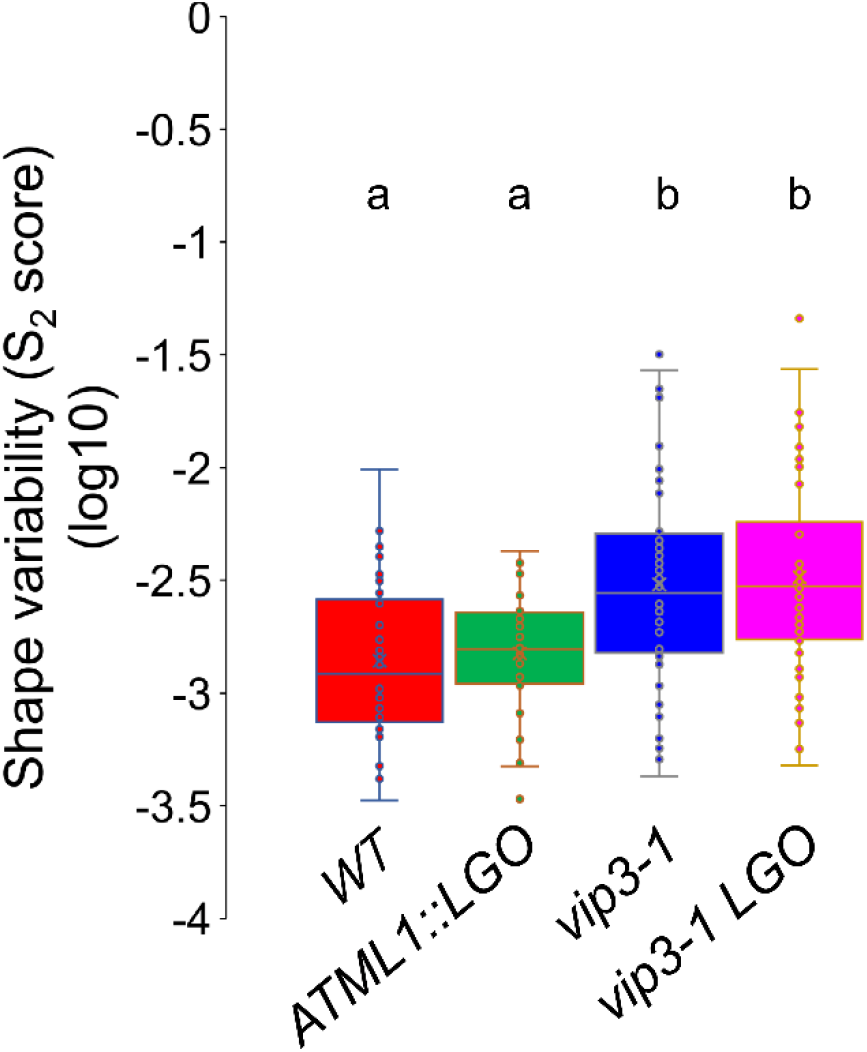
Sepal shape variability quantification in the wild type and lines with different mix of cell sizes Sepal shape variability is expressed as S_2_ score (squared deviation of sepal contours from the average contour) in log10 scale to aid with visualization. Higher score means higher shape variability. n = 40, 48, 50, 53 sepals for wild type, *pATML1::LGO (LGO), vip3-1* and *vip3-1 LGO*, respectively. 2-sided Welch’s t-test. Different letters indicate statistically significant differences (*p* ≤ 0.001).

The presence of giant cells could alter sepal shape in a couple of ways. First, giant cell precursors stop dividing early, so the number of cells or cell files across the sepal may be reduced, leading to a narrower shape. Second, because giant cells are usually long and straight, they can potentially influence the shape of the whole sepal, much like the ribs of a hand fan make the fan’s shape. Nevertheless, the effects of giant cells on the sepal shape in the wild-type background is quite small, probably because local growth, *i*.*e*. at the wall scale, is not majorly affected [17] and because global mechanisms channel growth pattern, for example, proper growth arrest at the sepal tip [5,6].

To further check the contribution of giant cells to sepal shape, we used a background where mechanisms for proper sepal growth are compromised. In the *vip3-1* mutant, gene expression becomes more variable, leading to increased variability in molecular growth regulators (ROS, auxins), increased local growth heterogeneity, and increased shape variability [6]. We reasoned that, in *vip3-1*, we might uncover stronger effects of giant cell overpopulation on final sepal shape. *vip3-1* sepals have a mixed of cell population, which is comparable to wild-type ones (Figure 1A). In wild-type sepals, the sepal tip stops growing early during sepal development, but that of *vip3-1* sepals keeps growing for longer [6]. To check our hypothesis, we introduced *pATML1::LGO* into the *vip3-1* background and measured sepal shape and shape variability (Figure 1A-C).

First, we extracted the average shape of *vip3-1* and *vip3-1 LGO* (*vip3-1 pATML1::LGO*) sepals and overlapped their average shapes (Figure 1D, right). *vip3-1* sepals were significantly wider than those of the wild type, as previously shown (Figure 1D-E; [6]). Regarding the contribution of giant cells to average sepal shape, we found that, as in the wild type, they make *vip3-1* sepals significantly narrower mostly at the lower half of the sepal (towards the base), leading to a slightly lower aspect ratio (0.46 for *vip3-1* and 0.43 for *vip3-1 LGO*; Figure 1D-E). To further understand this change, we measured the MWP index and found that while *pATML1::LGO* and *vip3-1* sepals have the MWPs similarly closer to the tip (MWP_*35S::LGO*_ = 0.40, MWP_*vip3-1*_ = 0.38, 5% difference), that of *vip3-1 LGO* is pushed significantly further to the tip (MWP_*vip3-1 LGO*_ = 0.30, 21% difference compared to MWP_*vip3-1*_) (Figure 2B). The large change in shape observed in *vip3-1 LGO* double mutant supports our hypothesis that the effects of giant cells would be exaggerated in a mutant with compromised mechanisms for organ growth.

We then calculated the score for sepal shape variability. Surprisingly, we found that there is no significant difference in shape variability between *vip3-1* and *vip3-1 LGO* sepals, i.e. similar to the comparison between wild-type and *pATML1::LGO* sepals (*S*_*2*_ score as median ± SE = 2.78 ± 1.0 10^−3^ for *vip3-1*, = 2.94 ± 1.1 10^−3^ for *vip3-1 LGO*) (Figure 3). This further confirms that sepal shape variability is robust to cell size heterogeneity. Note that since *vip3-1* already exhibits high shape variability, we cannot completely rule out the possibility that it is difficult to induce even higher variability by changing cell types.

Recently, another team was also independently investigating the roles of cell types in sepal shape robustness [22]. They showed that sepals having only giant cells (*pATML1::LGO*) or small cells (*lgo* mutant) have similar shape robustness compared to wild-type sepals [22], consistent with our results here. Using time-lapse imaging to analyze growth pattern at cellular level of wild-type, *pATML1::LGO* and *lgo* sepals, they associated their similar shape robustness with a similar cell growth pattern. When they introduced these lines into *ftsh4-5 (filamentous temperature sensitive H 4)*, a mutant with reduced shape robustness, they found that a population of small cells only (*lgo ftsh4-5*) significantly increase sepal shape variability in the *ftsh4-5* background, while giant cells (*pATML1::LGO ftsh4-5*) did not. The increase in sepal shape variability was associated with uneven growth rates and disorganized growth directions of cells [22]. Overall, their findings are complementary to ours, and provide a cell growth-based explanation for shape robustness of the mutants.

To summarize, our quantitative analyses reveal that: (i) a diverse cell population is not necessary for robust sepal shape, as demonstrated by similar shape variability between wild type and *pATML1::LGO*, (ii) while giant cells do not change shape variability, they could alter sepal shape in a subtle way, and (iii) in a background where growth variability is promoted and shape robustness is compromised (*vip3-1*), giant cells could induce more pronounced change in shape, but still did not affect shape variability. This suggests that organ shape variability does not emerge at the cell scale, but rather at smaller scales (e.g. individual cell wall properties, e.g. [17]) or larger scales (clones of cells, [23]). This means that the question of how organs know when to stop growing should be addressed with a multiscale lens.

## Data accessibility

The datasets analyzed during the current study, the code used to identify the Maximal width position in ImageJ, and the supplemental Table 1 are available from the OSF repository: https://osf.io/4aep5/ [24].

## Declaration of AI use

We have not used AI-assisted technologies in creating this article.

## Author contributions

D-C.T, C.T and O.H designed research; D-C.T conducted experiments and analyzed data; D-C.T prepared the original paper draft; C.L. wrote the ImageJ macro. All authors read, revised and approved the manuscript.

## Ackowledgements

We thank Dr. Adrienne Roeder (Cornell University, Ithaca, New York, USA) for *pATML1::LGO* seeds and for critical reading of the manuscript. We thank PLATIM platform for their help in using the Hirox microscope. This work is supported by the European Research Council (ERC, grant agreement No 101019515, “Musix”), by CEFIPRA (grant 6103-1), and by the French National Research Agency through a European ERA-NET Coordinating Action in Plant Sciences (ERA-CAPS) grant (Grant No. ANR-17-CAPS-0002-01).

